# Murine ex vivo cultured alveolar macrophages provide a novel tool to study tissue-resident macrophage behavior and function

**DOI:** 10.1101/2021.02.11.430791

**Authors:** A.-D. Gorki, D. Symmank, S. Zahalka, K. Lakovits, A. Hladik, B. Langer, B. Maurer, V. Sexl, R. Kain, S. Knapp

## Abstract

Tissue-resident macrophages are of vital importance as they preserve tissue homeostasis in all mammalian organs. Nevertheless, appropriate cell culture models are still limited. Here, we propose a novel culture model to study and expand murine primary alveolar macrophages (AMs), the tissue-resident macrophages of the lung, in vitro over several months. By providing a combination of GM-CSF, TGFβ and the PPARγ activator rosiglitazone, we maintain and expand mouse ex vivo cultured AMs, short mexAMs, over several months. MexAMs maintain typical morphologic features and stably express primary AM surface markers throughout in vitro culture. They respond to microbial ligands and exhibit an AM-like transcriptional profile, including the expression of AM specific transcription factors. Furthermore, when transferred into AM deficient mice, mexAMs efficiently engraft in the lung and fulfill key macrophage functions leading to a significantly reduced surfactant load in those mice. Altogether, mexAMs provide a novel, simple and versatile tool to study AM behavior in homeostasis and disease settings.

**KEYPOINTS:**

- A novel method to culture and expand primary alveolar macrophages over several months ex vivo
- Murine ex vivo cultured alveolar macrophages (mexAMs) restore lung function in a murine pulmonary alveolar proteinosis model

## INTRODUCTION

Tissue-resident macrophages (TRMs) are capable of self-renewal under homeostatic conditions in many organs including the lung^1,2^. By continuous sensing of the surrounding milieu, TRMs adapt to microenvironmental signals, resulting in distinct, tissue-specific macrophage identities^3–5^.

Alveolar macrophages (AMs), the TRMs of the lung, reside in the alveoli, the air-liquid interface of the lung, where they perform organ-specific functions such as the clearance of surfactant proteins and cell debris. AMs arise from fetal liver-derived monocytes that differentiate via granulocyte-macrophage colony-stimulating factor (GM-CSF)^1^ and transforming growth factor (TGFβ)^6^ induced expression of the transcription factor peroxisome proliferator-activated receptor gamma (PPARγ) around postnatal day 3^7,8^. The absence of autocrine TGFβ signaling in AMs of young mice resulted in reduced AM numbers, together with increased protein content in the bronchoalveolar lavage^6^. The accumulation of surfactant and subsequent development of pulmonary alveolar proteinosis (PAP) is also observed in mice and humans that lack mature AMs due to a loss of GM-CSF receptor subunits^9,10^. In patients, PAP is a rare lung disease associated increased susceptibility to infections and pulmonary fibrosis, which requires regular bronchoscopic removal of the protein-rich liquid^11,12^. To fully appreciate the functional versatility and therapeutic potential of AMs, appropriate cell culture models are required.

In vitro research on macrophages is essentially limited to the use of bone marrow-derived macrophages that can be expanded and cultured in sufficient numbers. In contrast, TRMs such as AMs must be isolated from mice and can only be kept in culture for a few days. In this study, we established a protocol for the ex vivo expansion and culture of primary AMs, which we termed mexAMs, by providing AMs with culture conditions that mimic lung microenvironmental factors. These cultured mexAMs expand rapidly and can be maintained and stored for several months, while continuously exhibiting characteristic features of primary AMs, including typical cell surface markers such as CD11c and Siglec-F and an AM-like transcriptional profile. Adoptively transferred mexAMs efficiently engraft in the lung and fulfill AM functions. This includes the reduction of surfactant and protein accumulation upon transfer into AM deficient mice. Taken together, mexAMs represent a valuable and versatile tool to study primary AM functions in health and disease.

## METHODS

### Mice

C57BL/6J, CD45.1^13^ and UBI-GFP^14^ mice were originally obtained from Jackson Laboratory. *CD169^Cre/+^STAT5ab^fl/fl^* mice (STAT5ΔCD169) or *STAT5^fl/fl^* littermate controls were obtained by crossing CD169-Cre^15^ provided by the RIKEN BRC, Japan, and floxed STAT5ab^16^ mice provided by R. Morrigl (University of Veterinary Medicine, Vienna). *Csf2rb^−/−^Csf2rb2^−/−^* mice^10,17^ were provided by M. Busslinger (Research Institute of Molecular Pathology, Vienna). All mice were maintained on a C57BL/6 background and bred and housed under specific pathogen-free conditions. Mice of both sexes, aged 7-14 weeks were used. All animal experiments were approved by the Austrian Federal Ministry of Sciences and Research (BMBWF-66.009/0340-V/3b/2019).

### Culture of murine postnatal liver cells

Livers were taken between postnatal day 0.5 and 5 and squeezed through a sterile 70 μm cell strainer. Red blood cells were lysed using ammonium chloride lysis buffer. Single cell suspensions (0.2×10^6^/ml) were cultured in RPMI 1640 containing 10% FCS and 1% pen/strep (“experiment medium”) supplemented with indicated combinations of murine GM-CSF (30 ng/ml), human TGFβ (10 ng/ml) and rosiglitazone (1 μM). After 6 days, adherent cells were detached and replated at 4×10^4^ cells/cm^2^. Subsequently, cells were passaged when 70-90% confluency was reached.

### Maintenance of mouse ex-vivo cultured alveolar macrophages (mexAM)

Bronchoalveolar lavage from single or a pool of three to four mice was used. Cells were counted and 0.2×10^6^ cells/ml were cultured in experiment medium supplemented with murine GM-CSF (30 ng/ml), human TGFβ (10 ng/ml) and rosiglitazone (1 μM). After 2 h non-adherent cells were washed off and fresh medium was added. Cells were passaged every 5-6 days when 70-90% confluency was reached.

### Isolation of murine bone marrow-derived macrophages and peritoneal macrophages

Femurs and tibias were flushed with PBS and mouse bone marrow cells were differentiated for 5 days in experiment medium supplemented with 10% L929 conditioned medium. Peritoneal lavage was performed using sterile PBS and cells were plated in experiment medium for 3 h and then washed twice to remove non-adherent cells. Peritoneal macrophages were immediately used for flow cytometry or RNA-seq protocols.

### Flow cytometry

Single cell suspensions were incubated with anti-mouse CD16/CD32 monoclonal antibody for 10 min at 4°C. A mix of fluorescently labeled monoclonal antibodies was added for 30 min at 4°C (Resource table). Sample acquisition was performed on a LSR Fortessa equipped with FACSDiva software (BD Biosciences). Singlets were gated using FSC-A versus FSC-H, followed by a FSC-A/SSC-A gate. Dead cells and erythrocytes were removed from analysis, using a fixable viability dye eFluor780 and anti-mouse Ter119. Next, CD45.1 and CD45.2 positive cells were used for further analysis. Alveolar macrophages were defined as SiglecF^high^CD11c^+^ cells.

### Seahorse measurement

Extracellular acidification rate (ECAR) and oxygen consumption rate (OCR) were analysed using a XF-96 Extracellular Flux Analyzer (Seahorse Bioscience) according to manufacturer’s instructions. Cells were plated in XF-96 cell culture plates (1×10^5 cells/well). To remove non-adherent cells, cells were washed twice with Seahorse XF RPMI medium supplemented with 10 mM Glucose, 1 mM Pyruvate, 2mM L-Glutamine and 3% FCS. For real-time analysis of ECAR and OCR, cells were pre-incubated for 1 h under non-CO_2_ conditions. To assess mitochondrial function, 1 μM oligomycin, 1.5 μM fluoro-carbonyl cyanide phenylhydrazone (FCCP) and 100 nM rotenone plus 1 μM antimycin A (all Sigma-Aldrich) were injected where indicated. Data analysis was performed on obtained OCR values after subtraction of respective non-mitochondrial respiration values from all data points. ATP production was represented as OCR difference to baseline after injection of oligomycin.

### Cytospin

Cells were spun onto glass slides using the Shandon Cytospin 4 and air-dried. Staining was performed using Giemsa solution.

### Tissue sampling and processing

Mice were sacrificed by isoflurane inhalation (3.5% isoflurane, 2 l/min oxygen) followed by intraperitoneal injection of 450 mg/kg ketamine and 37.5 mg/kg Rompun in sterile saline. A bronchoalveolar lavage with 1 ml saline was performed. Afterwards, lungs were mechanically disrupted by GentleMACS dissociation (Miltenyi Biotec) in RPMI 1640 containing 5% FCS, 165 U/ml Collagenase I and 12 U/ml Dnase I, followed by digestion for 30 min at 37°C and a final homogenization step. Cell suspension was passed through a 70 μm cell strainer and red blood cells were lysed using ammonium chloride buffer.

### Phagocytosis assay

Cells were plated for 3 h, followed by an incubation with FITC-labeled heat-inactivated *S. pneumoniae* (MOI 100) for 45 min at 37°C or 4°C (negative control). Uptake of bacteria was assessed via FITC expression using flow cytometry. Phagocytosis index was calculated as (MFI × % positive cells at 37°C) minus (MFI × % positive cells at 4°C).

### Electron microscopy

Cells were fixed in Karnovsky fixative (2% PFA, 2.5% GA in 0.1M cacodylate buffer), washed with cacodylate buffer and stored overnight at 4°C. Next, they were embedded in 1% agarose type IV and treated with 1% osmium tetroxide for 1 h and dehydrated through an ethanol series. After embedding in resin, ultrathin sections were placed on copper 150 mesh grids and stained with 2% uranyl acetate and lead citrate. Samples were examined with a JEM-1400 Plus transmission electron microscope (JEOL).

### Cell stimulations

Cell types were stimulated in 96-well plates (5×10^5^ cells/ml) with heat-inactivated *S. pneumoniae* (MOI 100) or LPS E.coli O55:B5 (10 ng/ml). The levels of secreted cytokines were determined in supernatants after 16 h. For polarization experiments cells were seeded for 2.5 h and afterwards treated with LPS (100 ng/ml) and IFNγ (200 U/ml), IL-4 (10 ng/ml) and IL-13 (10 ng/ml) or IL-10 (10 ng/ml) for 1.5 h (qPCR) or 16 h (nitrite measurement).

### Cytokine analysis

Indicated mouse cytokines were measured using the LEGENDplex Mouse Macrophage/Microglia Panel (BioLegend). Samples were prepared according to manufacturer's instructions and analysed by flow cytometry. Data analysis was performed using the LEGENDplex data analysis software.

### Proliferation assay

To assess cell proliferation, intracellular ATP levels were measured according to the CellTiter-Glo® assay (Promega) instructions. Luminescence was expressed as fold change compared to time-point of seeding.

### Immunocytochemistry

Lungs of untreated *Csf2rb^−/−^Csf2rb2^−/−^* mice or *Csf2rb^−/−^Csf2rb2^−/−^* that received GFP mexAMs intranasally 4 weeks earlier were fixed in 4% PFA for 48 h, dehydrated in 15% and 30% sucrose for 24 h each as described earlier^18^. Sections were stained with anti-pro + mature Surfactant Protein B antibody and DAPI.

### qPCR

Total mRNA was isolated using the NucleoSpin kit (Macherey-Nagel) according to manufacturers’ instructions. Real-time PCR was performed using the Perfecta SYBR Green Master Mix (Quant Bio). Following primers were used: m*Mrc1* (TCTGGGCCATGAGGCTTCTC, CACGCAGCGCTTGTGATC TT), m*Ym1* (TCTGGGTACAAGATCCCTGAACTG, GCTGCTCCATGGTCC TTCCA). Gene expression was normalized to m*Hprt* and expressed as fold change to indicated control.

### Nitrite measurement

Nitrite in cell culture supernatants was measured using the Griess Reagent System (Promega) according to manufacturers’ instructions.

### Adoptive cell transfer

Mice were anesthetized with isoflurane and 0.4-1×10^6^ cells per mouse were intranasally administered. Mice were sacrificed at indicated time-points. The bronchoalveolar lavage was centrifuged at 300 g and the optical density at 600 nm was recorded using an Ultrospec 10 (Amersham Biosciences). Cells were manually counted using a Neubauer chamber or a Z2 cell counter (Beckman Coulter).

### RNA-sequencing

Total RNA from indicated cell types was isolated using RNeasy Micro kit (Qiagen). Libraries were prepared from 150 ng total RNA input using the QuantSeq 3’ mRNA-Seq Library Prep Kit and UMI Second Strand Synthesis Module (Lexogen), according to the manufacturer’s instructions. Pooled libraries were 65□bp single-end sequenced on the HiSeq4000 (Illumina). Sequencing was performed at the Biomedical Sequencing Facility (CeMM and Medical University of Vienna). Demultiplexing of raw data and mapping to the mouse genome GRCm38 (mm10) was done using the Bluebee® software (version Quantseq 2.3.6 FWD UMI).

### DEG and GO enrichment

Differential expression analysis was performed using functions from the Bioconductor package DESeq2^19^. All macrophage populations derived from three independent biological replicates (mouse 1-3) or from two independent mexAM cultures taken at passage 8 and 13 (Pool 1) or passage 5 and 10 (Pool 2). DEGs were defined as absolute log2 fold change >2 and adjusted p-value <0.01 in comparisons between primary AMs and any other cell type (BMDM, PM, mexAM). Heatmaps were generated using the pheatmap function. The most significantly enriched GO terms were assessed using the enrichGO function of clusterprofiler^20^. The AM specific gene list was curated from two publications^3,21^ and unpublished data.

### Statistical analysis

Data are presented as mean ± SEM. Comparisons were performed using unpaired two-tailed Student’s *t*-test for two groups or one-way ANOVA for more than two groups. Statistical significance was defined as p< 0.05. Number of animals is indicated as “n”. Sizes of tested animal groups were dictated by availability of the transgenic strains and litter sizes, allowing littermate controls.

## RESULTS

### Murine alveolar macrophage-like cells can be derived from postnatal liver cells

To identify the optimal culture conditions for the expansion of AMs, we considered a previously published protocol, where murine fetal liver cells were cultured in the presence of GM-CSF^22^. Aiming for a setting that more closely resembles the lung microenvironment, we cultured postnatal murine liver cells in the presence of indicated combinations of GM-CSF, TGFβ and rosiglitazone, an activator of the AM transcription factor PPARγ^23^ (Fig. 1A). Already after six days, all cells treated with GM-CSF plus TGFβ or the triple combination had a round shape, closely resembling primary AMs (Fig. 1B, S1A). In contrast, two microscopically distinct cell populations, round and elongated, were observed when cells were grown in the presence of GM-CSF alone or GM-CSF plus rosiglitazone (Fig. 1B, S1A). In addition, we noticed that postnatal liver cells expanded very slowly in the presence of GM-CSF alone. This was reflected in significantly lower intracellular ATP levels in cells treated with GM-CSF alone (Fig. 1C, S1B).

**Fig 1.**
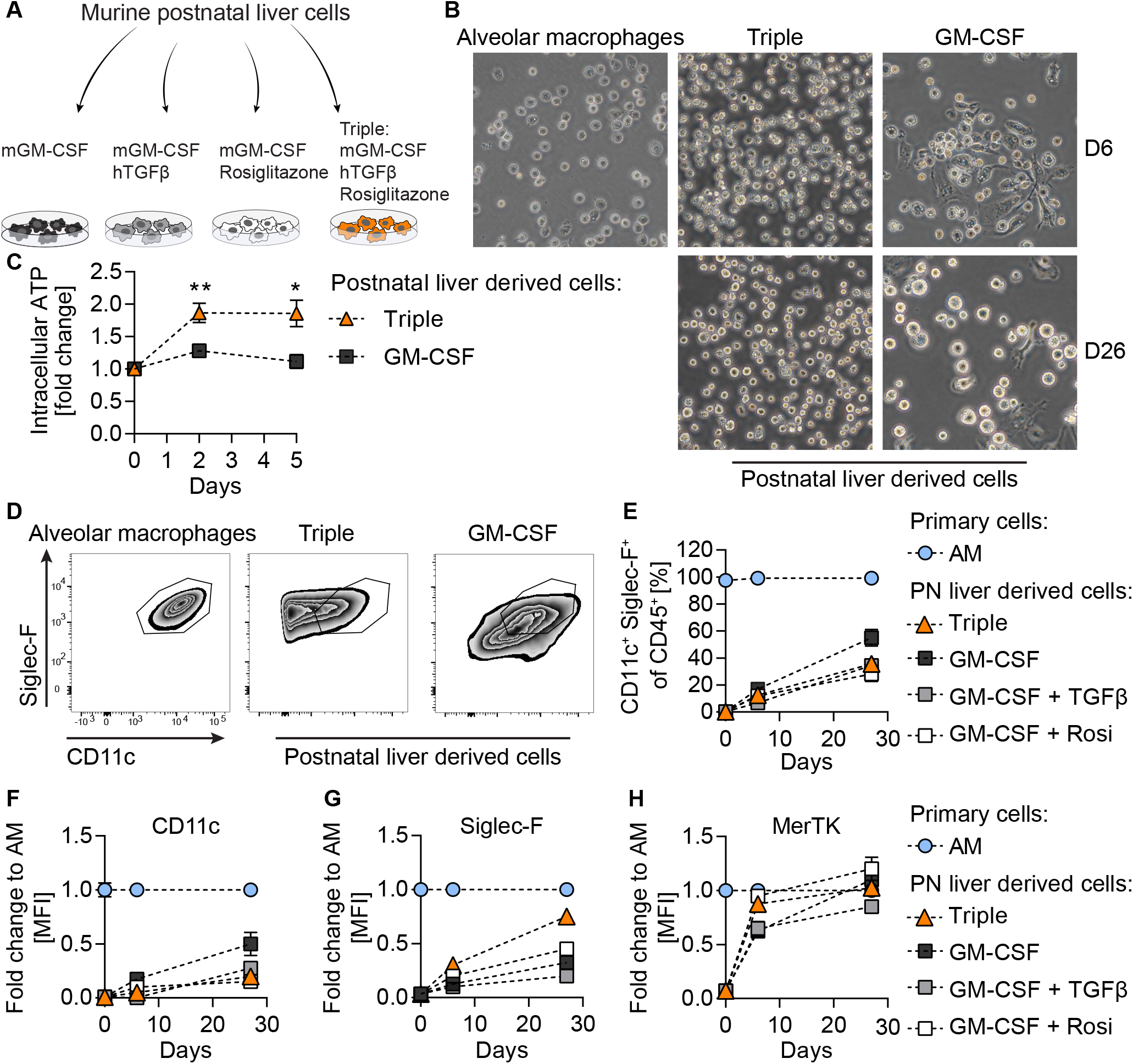
Murine alveolar macrophage-like cells can be derived from postnatal liver cells. (**A**) Experimental set-up. (**B**) Primary AMs after 3h in culture and postnatal liver cells treated with murine GM-CSF (30ng/ml) or murine GM-CSF (30ng/ml) + human TGFβ (10ng/ml) + rosiglitazone (1μM) (Triple) after 6 days (D6) and 26 days (D26) in culture; 40x magnification. (**C**) Cell proliferation of postnatal liver cells under indicated conditions over 5 days compared to time of seeding. (**D**) FACS analysis of Siglec-F and CD11c expression on primary AMs and postnatal liver cells grown under indicated conditions on D26. (**E**) Percentage of CD11c^+^Siglec-F^+^ cells in postnatal liver cell cultures at day of seeding, D6 and D26. (**F**) CD11c, (**G**) Siglec-F and (**H**) MerTK mean fluorescence intensity levels of postnatal liver cell cultures as fold change to primary AMs at indicated days. (**D**-**H**) Pre-gated on single, viable CD45^+^ cells. Graphs show means ± SEM of 3-4 biological replicates. Data are representative of at least two independent experiments. *p < 0.05, **p<0.01 (Student’s t test). AM= alveolar macrophages, D= day, PN= postnatal, Rosi= Rosiglitazone.

Next, we assessed the expression of AM specific as well as pan-macrophage surface markers to define the differentiation profile by flow cytometry. AMs typically express high levels of Siglec-F and CD11c^24^ (Fig. 1D, S1C). From day six onwards, CD11c^+^ Siglec-F^+^ cells emerged in the fetal liver cell cultures and the relative proportion of AM-like cells in culture gradually increased over time in all conditions (Fig. 1E, S1C). While CD11c was not expressed on freshly isolated postnatal liver cells, it increased over time, being highest on cells treated with GM-CSF only (Fig. 1F). Siglec-F expression was upregulated within 26 days in all conditions (Fig. 1G). Of note, cells treated with the combination of GM-CSF, TGFβ and rosiglitazone reached 100% of primary AM Siglec-F expression levels by differentiation day 77 (Fig. S1D, S1E). Mer tyrosine kinase (MerTK), a receptor involved in the engulfment of apoptotic cells, is highly expressed on various macrophage populations including AMs^25^. Already on differentiation day 6, MerTK expression was comparable to primary AM levels when cells were treated with GM-CSF plus rosiglitazone or the triple combination (Fig. 1H). As AMs develop from Ly-6C^+^CD11b^+^ fetal liver monocytes, we analyzed the expression of CD11b and Ly-6C over time, and observed a gradual downregulation of Ly-6C and consistently very low levels of CD11b (Fig. S1F, S1G).

These results show the ability of murine postnatal liver cells, cultured in the presence of GM-CSF, TGFβ and rosiglitazone, to progressively transition from a monocyte phenotype to an AM-like morphology and marker profile, while maintaining their proliferative capacity.

### Murine ex vivo cultured alveolar macrophages are functionally similar to primary alveolar macrophages

TRMs are terminally differentiated immune cell populations that retain self-renewing capacities^2,26,27^. Being able to generate AM-like cells from murine fetal liver cells, we continued to test our optimized protocol on mature, primary murine AMs (Fig. 2A). Primary AMs expanded and maintained their CD11c^+^ Siglec-F^+^ cell expression profile in culture over six months when treated with the combination of GM-CSF, TGFβ and rosiglitazone (Fig. 2B, S2A). These murine ex vivo cultured AMs (mexAMs) appeared strikingly similar to primary AMs, but distinct from BMDMs (Fig. 2C, S2B).

**Fig 2.**
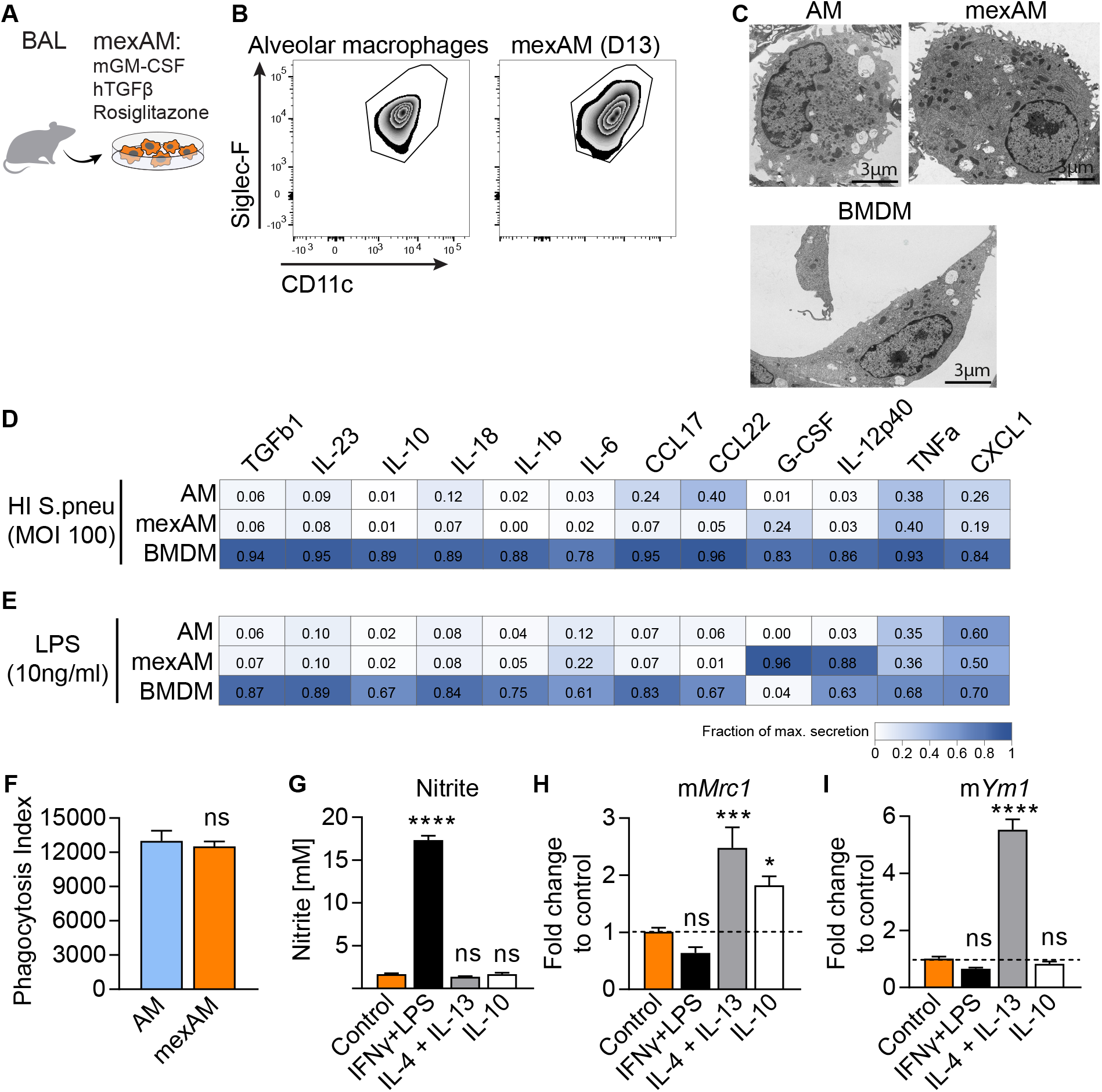
Murine ex vivo cultured alveolar macrophages (mexAMs) are functionally similar to primary alveolar macrophages. (**A**) Experimental set-up. (**B**) FACS analysis of Siglec-F and CD11c expression on primary AMs and mexAMs on day 13. Pre-gated on viable CD45^+^ cells. (**C**) Electron microscopy pictures of primary AMs, mexAMs or BMDMs. Magnification: 3000x, scale bar: 3μm. (**D** and **E**) Measurement of indicated cytokines upon heat-inactivated *S. pneumoniae* (MOI 100) (**D**) or LPS (10ng/ml) (**E**) stimulation of AMs, mexAMs or BMDMs for 16 h, expressed as fraction of maximal secretion. (**F**) Phagocytosis index of primary AMs and mexAMs. (**G**) Nitrite concentration in supernatants of polarized mexAMs after 16 h. (**H** and **I**) M2 polarization markers m*Mrc1* (**H**) and m*Ym1* (**I**) assessed by RT-PCR in polarized mexAMs after 1.5 h. Graphs show means ± SEM of 3-4 biological replicates (D, E) or technical quadruplicates (F-I). Data are representative of at least two independent experiments. *p < 0.05, **p<0.01, ***p<0.001, ****p<0.0001 (one-way ANOVA followed by Dunnett’s multiple comparison test). AM= alveolar macrophages, BAL= bronchoalveolar lavage, BMDM= bone marrow-derived macrophages, D= day, HI= heat-inactivated, mexAM= mouse ex vivo cultured alveolar macrophages.

As innate immune cells, macrophages play a key role in the defense against pathogens by initiating a pro-inflammatory response. To test their functional properties, we exposed mexAMs, AMs and BMDMs to heat-inactivated *S. pneumoniae* (HI *S. pneu*, Fig. 2D, Table S1) and lipopolysaccharide (LPS, Fig. 2E, Table S1). With a few exceptions, mexAMs responded like primary AMs and showed a less vigorous release of cytokines and chemokines than BMDMs (Fig. 2D, 2E). We also tested if a freeze-thaw cycle affects the responsiveness of mexAMs and discovered that IL-6 (Fig. S2C) and CXCL1 (Fig. S2D) levels did not differ between HI *S. pneu* stimulated thawed and continuously cultured mexAMs.

A main function of AMs in situ pertains to the phagocytosis of surfactant proteins and cellular debris in the alveoli. To test the phagocytic activity of mexAMs compared to primary AMs, we incubated them with FITC-labeled HI *S. pneu*, and observed an efficient and comparable uptake of bacteria by both cell types (Fig. 2F). Another key characteristic of macrophages is the plasticity in their response to stimuli they are exposed to, while constantly surveying the surrounding tissue^28^. To assess this, we polarized mexAMs with classically activating M1 (IFN-γ and LPS), alternatively activating M2 (IL-4 and IL-13), as well as deactivating (IL-10) stimuli. MexAMs maintained their plasticity, illustrated by the nitrite release upon M1-polarization (Fig. 2G) and induction of mannose receptor, C type I (*Mrc1*, Fig. 2H) and chitinase-like 3 (*Ym1*, Fig. 2I) upon M2-polarization. To investigate if mexAMs can be maintained without the combination of GM-CSF, TGFβ and rosiglitazone, we removed all trophic factors from the medium, which resulted in cell death of mexAMs within one week (Fig. S2E).

Collectively, these data demonstrate that mexAMs, in contrast to BMDMs, exhibit phenotypic and functional properties of primary AMs.

### MexAMs are phenotypically and transcriptionally closest to primary murine alveolar macrophages

Having established that mexAMs are functionally similar to primary AMs, we next compared their surface marker and transcriptional profile to different macrophage populations. These included BMDMs, as the macrophage type predominantly used for in vitro studies, arising from myeloid bone marrow progenitor cells, as well as peritoneal macrophages (PMs), a mature TRM population exposed to a different tissue microenvironment. Cell surface expression levels of the pan-macrophage marker F4/80 were comparable between all macrophage types, with slightly higher levels in mexAMs and PMs (Fig. 3A). In contrast, the AM surface markers Siglec-F (Fig. 3B) and CD11c (Fig. 3C) were exclusively expressed on mexAMs and AMs, but not BMDMs and PMs.

**Fig 3.**
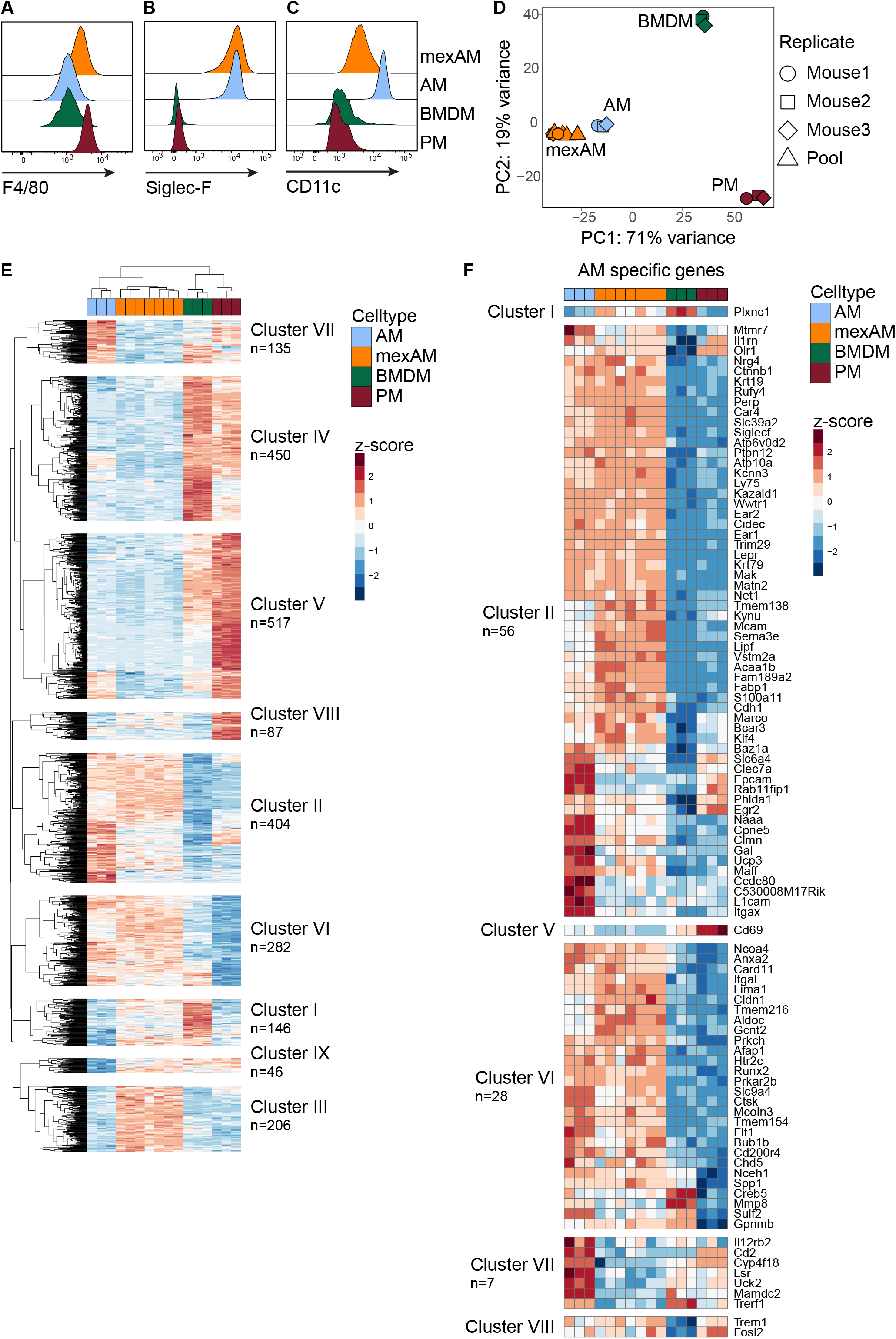
MexAMs are phenotypically and transcriptionally closest to primary murine alveolar macrophages. (**A**) F4/80, (**B**) Siglec-F and (**C**) CD11c cell surface expression on mexAMs, primary AMs, primary PMs and BMDMs measured by FACS. Pre-gated on viable CD45^+^ cells. (**D**) PCA analysis of indicated cell types of three biological replicates (mouse 1-3) plus mexAM samples derived from different passages of mexAM cultures (Pool). (**E**) Heatmap of genes differentially expressed (absolute log2fc value >2, p-adj <0.01) between primary AMs and any other cell type (PM, BMDM, mexAM). n indicates total number of genes per cluster. Raw counts were rlog transformed, followed by z-score scaling. (**F**) Heatmap of AM specific genes found in indicated clusters. AM= alveolar macrophages, BMDM= bone marrow-derived macrophages, mexAM= mouse ex vivo cultured alveolar macrophages, PC= principal component, PM= peritoneal macrophages.

By examining differences in the transcriptional profile of these four macrophage types, principal component analysis conclusively revealed that mexAMs and AMs clustered tightly together, whereas BMDMs and PMs showed distinct transcriptional profiles (Fig. 3D). Consistently, hierarchical clustering of 2273 differentially expressed genes (DEG) revealed that mexAM samples clustered next to AMs, whereas BMDM samples were closest to PMs (Fig. 3E). We identified nine different gene clusters (Fig. 3E, Table S2) with cluster IV and V consisting of genes upregulated in BMDMs and PMs. Cluster VIII comprised PM-specific genes such as *Klf2* and *Naip1*. The BMDM specific cluster I contained genes previously shown to be highly expressed in BMDMs such as *Trem2*^29^. Most interesting were two prominent clusters of genes, cluster II and VI, because they were upregulated in AMs and mexAMs but downregulated in BMDMs and PMs. When we compiled a list of 133 AM specific transcription factors and genes^3,21^ (Table S3), we found 97 to be included in the DEG list shown in Fig. 3E. Most of these genes including *Ear2*, *Marco*, *Fabp1* as well as the transcription factor *Klf4* were part of the two clusters (II and VI) shared between mexAMs and AMs (Fig. 3F). Similarly, the transcription factor *Car4*, which is uniquely expressed in AMs^4^, was elevated in all mexAM and AM samples (cluster II). *Itgax*, the gene underlying CD11c, was highly upregulated in AMs but less expressed on a transcriptional level in mexAMs, whereas *Siglecf* was highly expressed in mexAM and AM samples, coinciding with the flow cytometry results shown before (Fig. 3B, 3C). In the mexAM specific cluster (cluster III), many metabolic genes such as *Acly*, *Pdk1* or *Fasn* were upregulated (Fig. S3A). This was also reflected in the enriched GO terms, which included different metabolic processes (Fig. S3B). Seahorse experiments confirmed highly elevated basal OCR, ATP production and basal ECAR levels in actively expanding mexAMs when compared to primary AMs and terminally differentiated BMDMs (Fig. S3C-G).

These data led us to conclude that mexAMs and primary AMs share a common transcriptional signature that differs from BMDMs and PMs.

### MexAMs engraft efficiently in a partially depleted alveolar macrophage niche in vivo

To test whether mexAMs engraft in the physiological AM niche in vivo, we transferred CD45.1^+^ mexAMs by intranasal administration into CD45.2 expressing STAT5ΔCD169 mice or littermate controls (Fig. 4A). STAT5 is required for the development of lung dendritic cells (DC) and AMs^30^. The loss of STAT5 in CD169 expressing cells, which include AMs but not monocytes or DCs^31^, led to a partially emptied AM niche, indicated by a significantly reduced number of AMs in the bronchoalveolar lavage fluid (BALF) (Fig. S4A). Four weeks after transfer, a pronounced CD45.1^+^ mexAM population was found in the BALF of STAT5ΔCD169 and control mice (Fig. 4B, upper and lower right panel). In STAT5ΔCD169 mice we observed up to 50% of CD11c^+^Siglec-F^+^ cells to be of CD45.1^+^ mexAM origin, while in littermate controls about 9% of BALF (Fig. 4C) or lung (Fig. S4B) CD11c^+^Siglec-F^+^ cells expressed CD45.1. When expressed in absolute numbers, mexAM transfer sufficiently restored the AM niche in STAT5ΔCD169 animals after four weeks (Fig. 4D).

**Fig 4.**
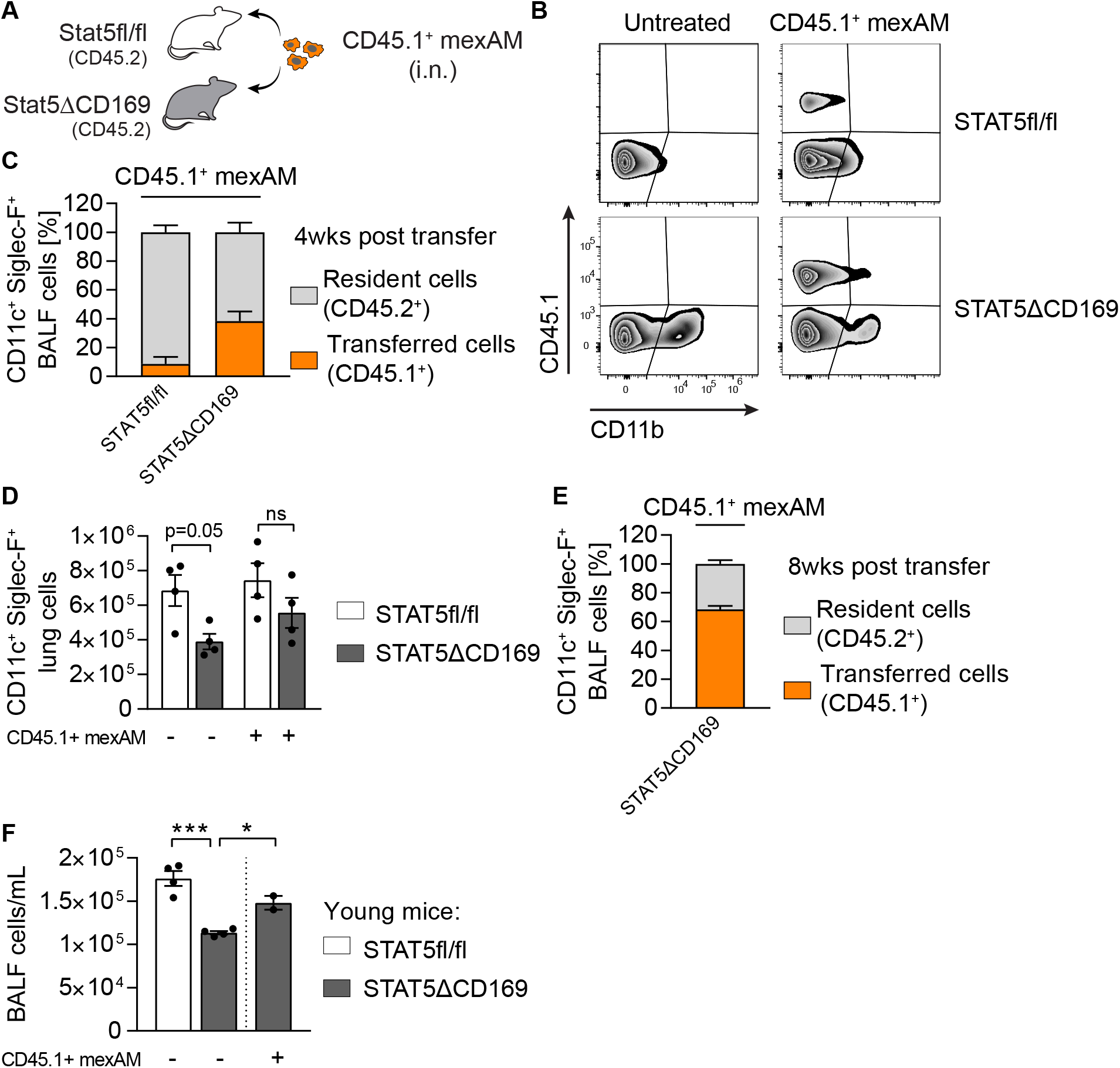
MexAMs engraft efficiently in a partially depleted alveolar macrophage niche in vivo. (**A**) Experimental set-up. Intranasal transfer of CD45.1^+^ mexAMs into CD45.2 expressing control (STAT5fl/fl) or STAT5ΔCD169 mice. (**B**) FACS analysis of CD45.1 and CD11b expression of control (upper panel) and STAT5ΔCD169 (lower panel) BALF cells untreated, or 4 weeks after transfer of CD45.1^+^ mexAMs. Pregated on viable Siglec-F^+^/CD11c^+^ cells. (**C**) Percentage of resident (CD45.2^+^, grey) and transferred (CD45.1^+^, orange) cells in BALF of control (STAT5fl/fl, n=4) and STAT5ΔCD169 mice (n=4) 4 weeks post CD45.1^+^ mexAM transfer. (**D**) CD11c^+^Siglec-F^+^ lung cell number in STAT5fl/fl and STAT5ΔCD169 mice untreated and 4 weeks post transfer of CD45.1^+^ mexAMs. (**E**) Percentage of resident (CD45.2^+^, grey) and transferred (CD45.1^+^, orange) cells in BALF of STAT5ΔCD169 mice (n=4-5) 8 weeks post CD45.1^+^ mexAM transfer. (**F**) BALF cell count per ml in STAT5fl/fl and STAT5ΔCD169 mice 14 weeks post transfer of CD45.1^+^ mexAMs into young (14 d) old mice. Graphs show means ± SEM of 2-5 biological replicates. *p < 0.05, **p<0.01, ***p<0.001 (one-way ANOVA followed by Sidak’s multiple comparison test). BALF= bronchoalveolar lavage, i.n.= intranasal, mexAM= mouse ex vivo cultured alveolar macrophages, ns= non significant.

To test if transferred mexAMs are capable to long-term repopulate the alveolar niche^2^, we transferred CD45.1^+^ mexAMs into the partially empty AM niche of STAT5ΔCD169 mice and analyzed the BALF eight weeks later. We found a prominent CD45.1^+^ mexAM population, comprising up to 73% of the total AM population in STAT5ΔCD169 mice. This indicates that transferred mexAMs self-renew and repopulate the AM niche long-lasting (Fig. 4E, S4C). Finally, we tested if mexAMs possess the ability to settle in and repopulate the lungs of newborn mice. Thus, we transferred CD45.1^+^ mexAMs into two weeks old STAT5ΔCD169 animals. Consistent with the results in adult STAT5ΔCD169 mice, we found a significant increase in BALF cell numbers 14 weeks after mexAM transfer into young mice (Fig. 4F). These findings demonstrate that mexAMs can efficiently engraft and replenish the AM niche in vivo.

### MexAMs restore lung function in a murine pulmonary alveolar proteinosis model

To understand if mexAMs can home to the AM niche as efficiently as primary AMs, we transferred CD45.1^+^ mexAMs and GFP^+^ AMs in a 1:1 ratio into the partially depleted AM niche of STAT5ΔCD169 mice (Fig. 5A). Using flow cytometry we could clearly distinguish these two populations after the transfer (Fig. 5B) and on average 27% of BALF cells consisted of transferred cells after four weeks (Fig. S5A). Analysis of the transferred CD11c and Siglec-F expressing population revealed that the ratio between CD45.1^+^ mexAMs and GFP^+^ primary AMs remained unchanged and that mexAMs and primary AMs contributed equally to the AM population (Fig. 5C). Twelve weeks after transfer, BALF cells in STAT5ΔCD169 mice mainly comprised of transferred cells (Fig. S5C), consistent with data shown in Fig. 4E. However, at this time GFP^+^ primary AMs outnumbered CD45.1^+^ mexAMs (Fig. S5B). To compare the proliferation and homing potential of two in vitro cultured macrophage types, we repeated the experimental set-up described in Fig. 5A and transferred GFP^+^ BMDMs and CD45.1^+^ mexAMs in a 1:1 ratio (Fig. S5D). Already after four weeks, BMDMs made up a higher proportion of the transferred cells in the immune cell population of the BALF (Fig. S5E-F).

**Fig 5.**
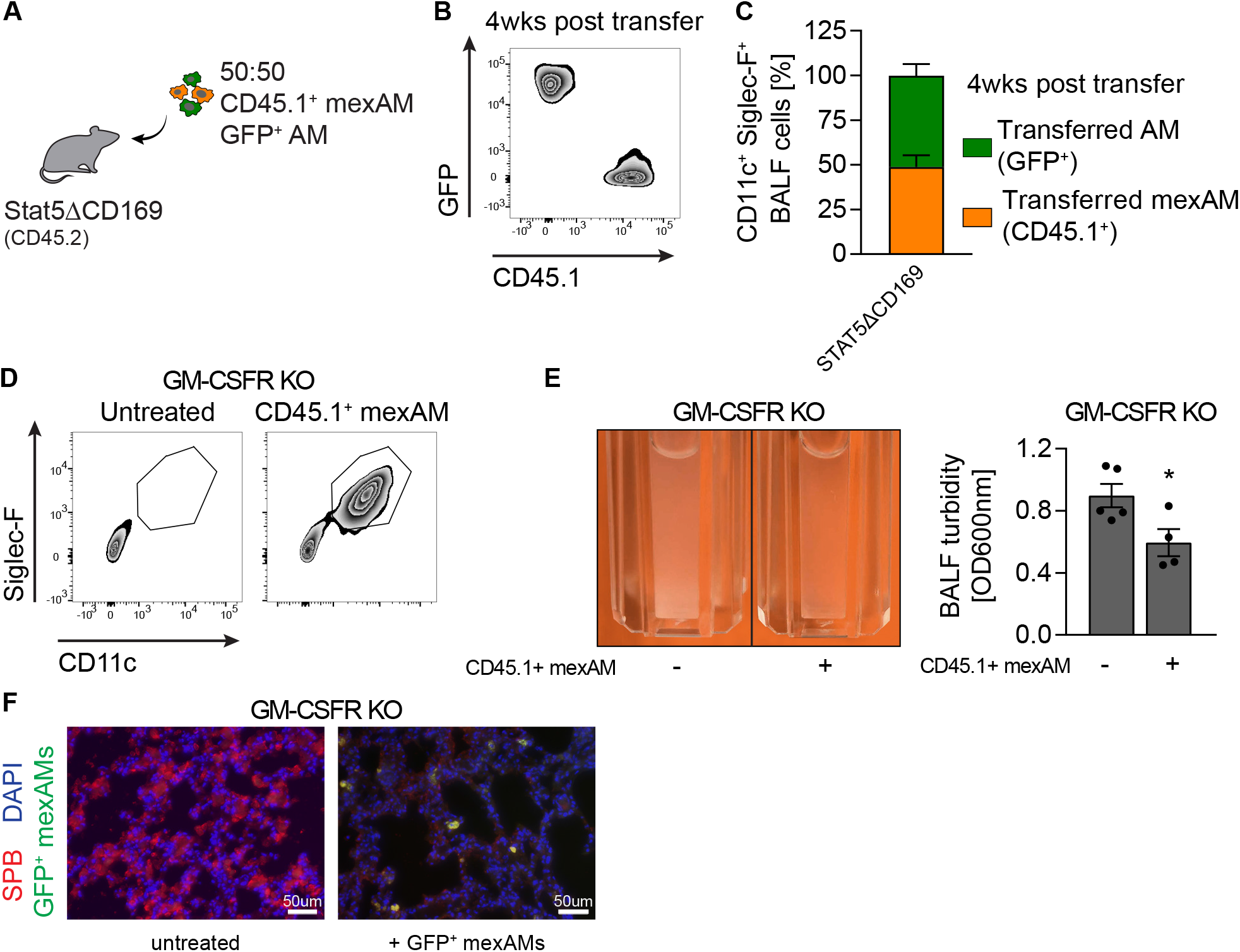
MexAMs restore lung function in a murine pulmonary alveolar proteinosis model. (**A**) Experimental set-up. Intranasal transfer of CD45.1^+^ mexAMs and GFP^+^ AMs in a 1:1 ratio into CD45.2 expressing STAT5ΔCD169 mice. (**B**) FACS analysis of GFP and CD45.1 expression in BALF of STAT5ΔCD169 mice 4 weeks after transfer of 50% GFP^+^ AMs and 50% CD45.1^+^ mexAMs. Pre-gated on viable Siglec-F^+^/CD11c^+^ cells. (**C**) Percentage of AMs (GFP^+^, green) and mexAMs (CD45.1^+^, orange) Siglec-F^+^/CD11c^+^ cells in BALF of STAT5ΔCD169 (n=4) mice 4 weeks post transfer. (**D**) FACS analysis of Siglec-F and CD11c expression on cells in BALF of GM-CSF receptor knock-out mice with and without transferred CD45.1^+^ mexAMs. (**E**) Representative picture and quantification of BALF turbidity of GM-CSF receptor knock-out mice control and after transfer of CD45.1^+^ mexAMs (4 weeks). (**F**) Immunofluorescent picture of surfactant protein B accumulation (SFB, red) in GM-CSF receptor knock-out mice without (left) and 4 weeks after transfer of GFP^+^ mexAMs (green). Magnification: 20x, scalebar: 50 μm. Graphs show means ± SEM of 4-5 biological replicates or representative pictures. *p < 0.05 (Student’s t test). BALF= bronchoalveolar lavage, mexAM= mouse ex vivo cultured alveolar macrophages.

One of the essential housekeeping functions of AMs is the clearance of lipids and proteins from the alveolar space^32^. Impaired GM-CSF signaling leads to the absence of mature and functional AMs, increased accumulation of surfactant proteins and subsequent development of pulmonary alveolar proteinosis (PAP)^12^. To assess if mexAMs are capable of lipid and protein clearance, we transferred CD45.1^+^ mexAMs into lungs of GM-CSF receptor knock out mice (*Csf2rb^−/−^Csf2rb2^−/−^*, GM-CSFR KO). Four weeks thereafter, we detected a CD11c^+^ Siglec-F^+^ AM population in the lavage (Fig. 5D). Development of PAP and surfactant accumulation was significantly reduced upon mexAM transfer, as illustrated by a significantly reduced BALF turbidity (Fig. 5E), and a decreased surfactant protein B content in the lungs of GM-CSFR KO mice (Fig. 5F).

These data show that mexAMs engraft the alveolar niche and take over AM functions in vivo, thereby preventing PAP development in GM-CSFR KO mice. Collectively, these data support the usefulness of mexAMs to study AM behavior in homeostatic and disease settings.

## DISCUSSION

The tissue environment shapes the identity of macrophage subsets^3–5^, making it almost mandatory to use specific, tissue derived macrophages for in vitro studies. Furthermore, most TRM populations are of embryonic origin^2^ and are thereby quite different in their development compared to the widely used, adult hematopoietic stem cell derived BMDMs.

By mimicking the lung microenvironment using GM-CSF, TGFβ and the PPARγ activator rosiglitazone we first used fetal liver cells to generate AM-like cells. Supplementing the medium with GM-CSF induced expression of the integrin molecule CD11c in accordance with established protocols that use GM-CSF to generate CD11c expressing DCs from hematopoietic progenitors^33,34^. The addition of TGFβ to the culture medium induced constant expansion of fetal liver derived AM-like cells, confirming the importance of TGFβ in the early AM differentiation^6^. Notably, only the use of the combination of GM-CSF, TGFβ and rosiglitazone induced high Siglec-F expression levels, a hallmark of primary AMs. We propose to use these fetal liver-derived AM-like cells as a tool to study AM development in vitro.

In a next step we cultured mature, fully differentiated primary AMs ex vivo. MexAMs expand rapidly while keeping their phenotypic and functional AM-like profile over several months in culture. In line with published results^1,8^ we found that expansion of mexAMs is dependent on GM-CSF and cannot be accomplished by supplementing medium with TGFβ or rosiglitazone alone (data not shown). While limited GM-CSF concentrations regulate the population of the AM niche in vivo^23^, excessive GM-CSF concentrations used under culture conditions allow constant mexAM proliferation and expansion.

A main advantage of any in vitro culture system is the unrestricted number of cells that can be utilized for high-throughput assays like whole genome CRISPR screens. We used a common transfection reagent and could confirm that mexAMs can be efficiently transfected with established protocols (data not shown), supporting their usability for high-throughput screening experiments.

Analysis of the transcriptional profile of mexAMs revealed that these cells clustered most closely with AM samples and maintained an AM specific gene expression profile including the transcription factors Klf4^35^ and Car4^4^. The fact that these mexAM samples derived from single biological replicates or pooled lavages as well as from different passaging numbers, confirms the reproducibility and stability of the AM transcriptomic profile over time. Yet, it is important to note that mexAMs, as an actively proliferating population, exhibited features of expanding cells in their metabolic profile. We therefore recommend additional optimization strategies, when using mexAMs for metabolic assays.

In the next step, we demonstrated that transferred mexAMs replenished the partially depleted AM niche of STAT5ΔCD169 mice and could be detected up to 14 weeks later in the lavage. Remarkably, even when transferred to WT mice mexAMs settled in the filled niche, albeit to a much lower extent than in STAT5ΔCD169 mice, supporting the idea that macrophage niches are self-regulating systems that contain a stable macrophage number^23^. When performing competitive transfers with 1:1 ratio of mexAMs and AMs, we detected equal numbers of both populations after four weeks. Surprisingly though, when we repeated the competitive transfer assay, mixing mexAMs and BMDMs, we saw a higher proliferative capacity of BMDMs as compared to mexAMs. The high proliferative capacity of transferred BMDMs is in line with data showing that adult bone marrow monocytes display a competitive advantage when re-filling an emptied AM niche, as compared to the remaining donor derived AM population^1,2,36^.

We envision a wide range of potential applications for mexAMs including co-culture models with immune and structural cells, to better understand disease settings associated with macrophages including lung fibrosis^36^ or cancer^37^. Here, we provide evidence that mexAMs can substitute for primary AMs as they restored impaired alveolar surfactant cleaning and prevented the development of PAP in mice lacking primary AMs in vivo.

In summary, our study highlights a previously underappreciated ability to culture and expand fully differentiated TRMs ex vivo over several months by maintaining their intrinsic, tissue-resident macrophage profile.

## Supporting information

Resource table

## ACKNOWLEDGMENTS

We thank the flow cytometry core facility and the animal facility of the Medical University of Vienna for their support. CD169cre mice were kindly provided by Miriam Merad (Icahn School of Medicine at Mount Sinai, New York, USA). S.K. is supported by the Austrian Science Fund (FWF) within the Special Research Programs Chromatin Landscapes (L-Mac: F 6104) and Immunothrombosis (F 5410), as well as the Doctoral Program Cell Communication in Health and Disease (W1205). V.S. and B.M. are supported by the FWF Special Research Program (F 6107).

## AUTHOR CONTRIBUTIONS

A.-D.G., D.S., S.Z., K.L. and A.H. performed experiments and analyzed data; A.-D.G. analyzed bioinformatic data; B.L. and R.K. performed electron microscopy experiments; B.M. and V.S. provided valuable reagents and technical advice. A.-D.G. and S.K. wrote the manuscript with input from co-authors and conceptualized the study.

## Supplemental figure legends

**Fig. S1.**
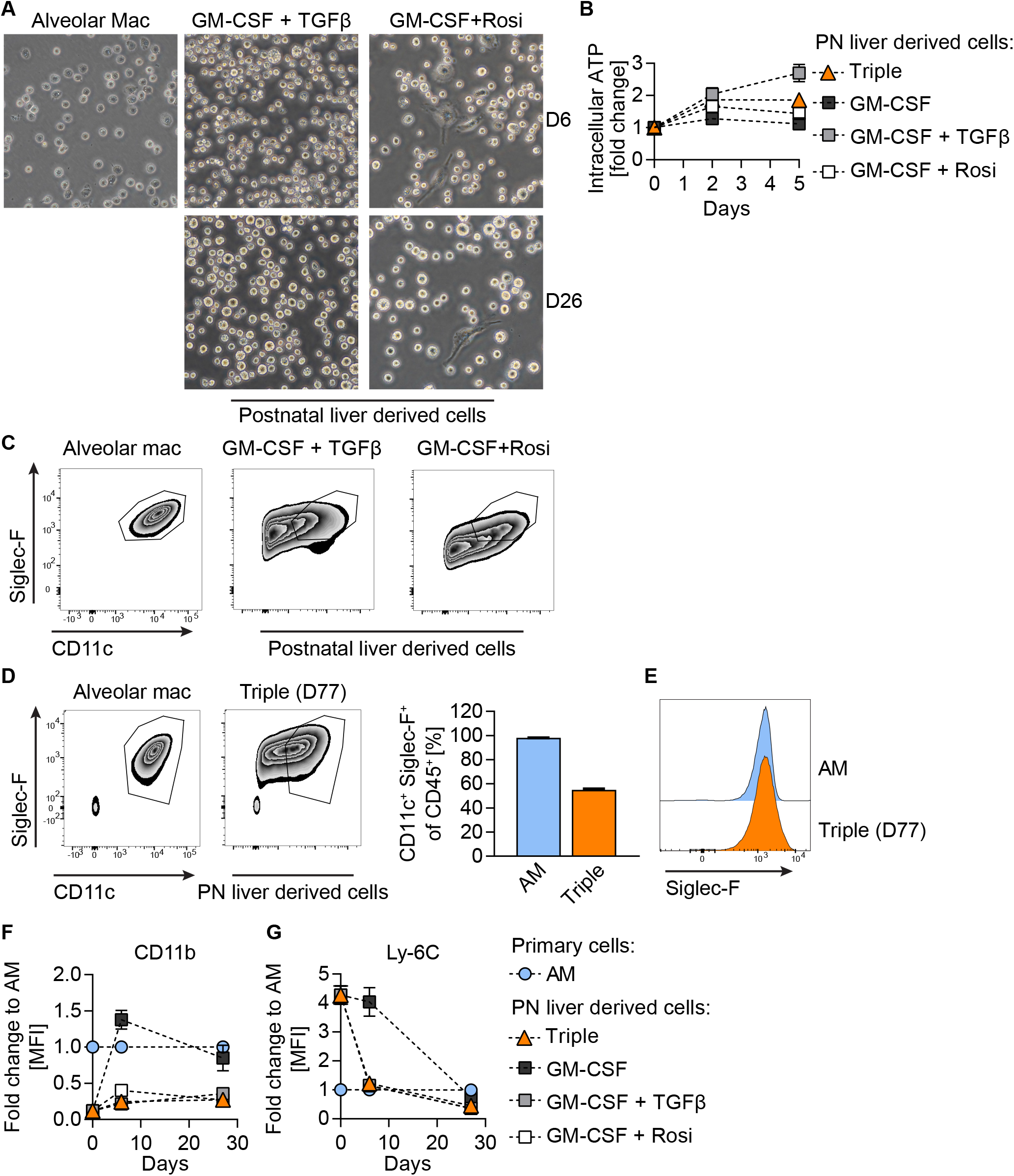
(**A**) Primary AMs after 3 h in culture and postnatal liver cells treated with murine GM-CSF (30ng/ml)+ human TGFβ (10ng/ml) or GM-CSF (30ng/ml)+ Rosiglitazone (1μM) after 6 d (D6) and 26 d (D26) in culture; 40x magnification. (**B**) Cell proliferation of postnatal liver cells under indicated conditions over 5 days compared to time of seeding. (**C**) FACS analysis of Siglec-F and CD11c expression in primary AMs and postnatal liver cells grown under indicated conditions on D26. (**D**) FACS analysis of Siglec-F and CD11c expression in primary AMs and postnatal liver cells grown with murine GM-CSF+ human TGFβ+ Rosiglitazone (Triple) on D77. (**E**) Siglec-F expression in primary AMs and postnatal liver cells grown with murine GM-CSF+ human TGFβ+ Rosiglitazone (triple) on D77. (**F**) CD11b and (**G**) Ly-6C mean fluorescence intensity levels of postnatal liver cell cultures as fold change to primary alveolar macrophages at indicated days. (**C-G**) Pre-gated on single, viable CD45^+^ cells. Graphs show means ± SEM of 3-4 biological replicates. Data are representative of at least two independent experiments. AM= alveolar macrophages, D= day, PN= postnatal, Rosi= Rosiglitazone.

**Fig. S2.**
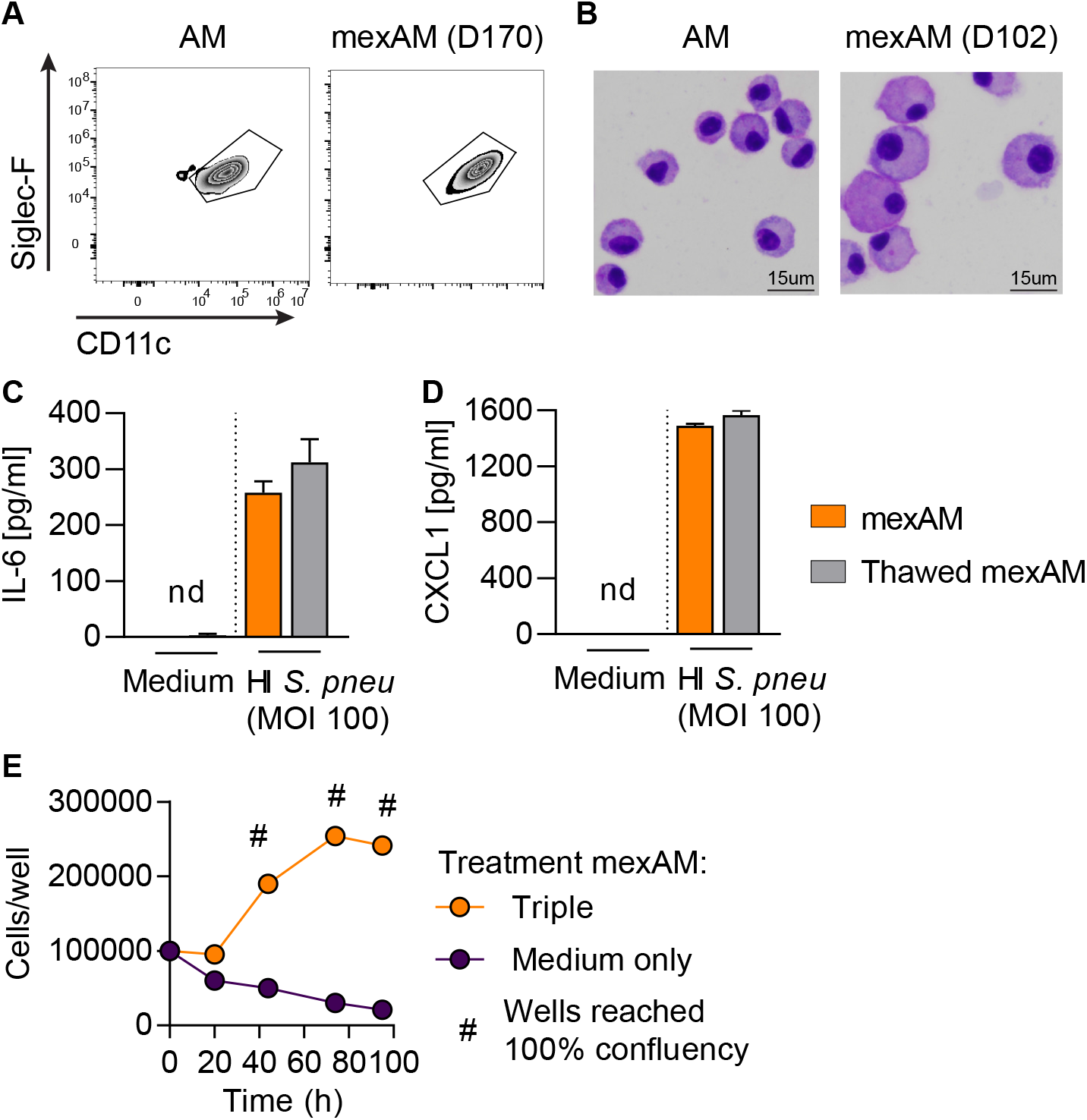
(**A**) FACS analysis of Siglec-F and CD11c expression in primary AMs and mexAMs after 170 days in culture. Pre-gated on viable CD45^+^ cells. (**B**) Cytospin pictures of Giemsa-stained primary AMs or mexAMs cultured for 102 days. Magnification: 40x, scale bar: 15μm. (**C** and **D**) IL-6 (**C**) and CXCL-1 (**D**) levels in supernatants of continuously cultured and thawed mexAMs stimulated with heat-inactivated *S. pneumoniae* (MOI 100) for 16 h. (**E**) Cell number of mexAMs cultured in murine GM-CSF+ human TGFβ+Rosiglitazone (Triple) containing medium or in medium without trophic factors over time. Graphs show means ± SEM of technical quadruplicates. Data are representative of at least two independent experiments. AM= alveolar macrophages, D= day, h= hours, mexAM= mouse ex vivo cultured alveolar macrophages, nd= non detectable, MOI= multiplicity of infection.

**Fig. S3.**
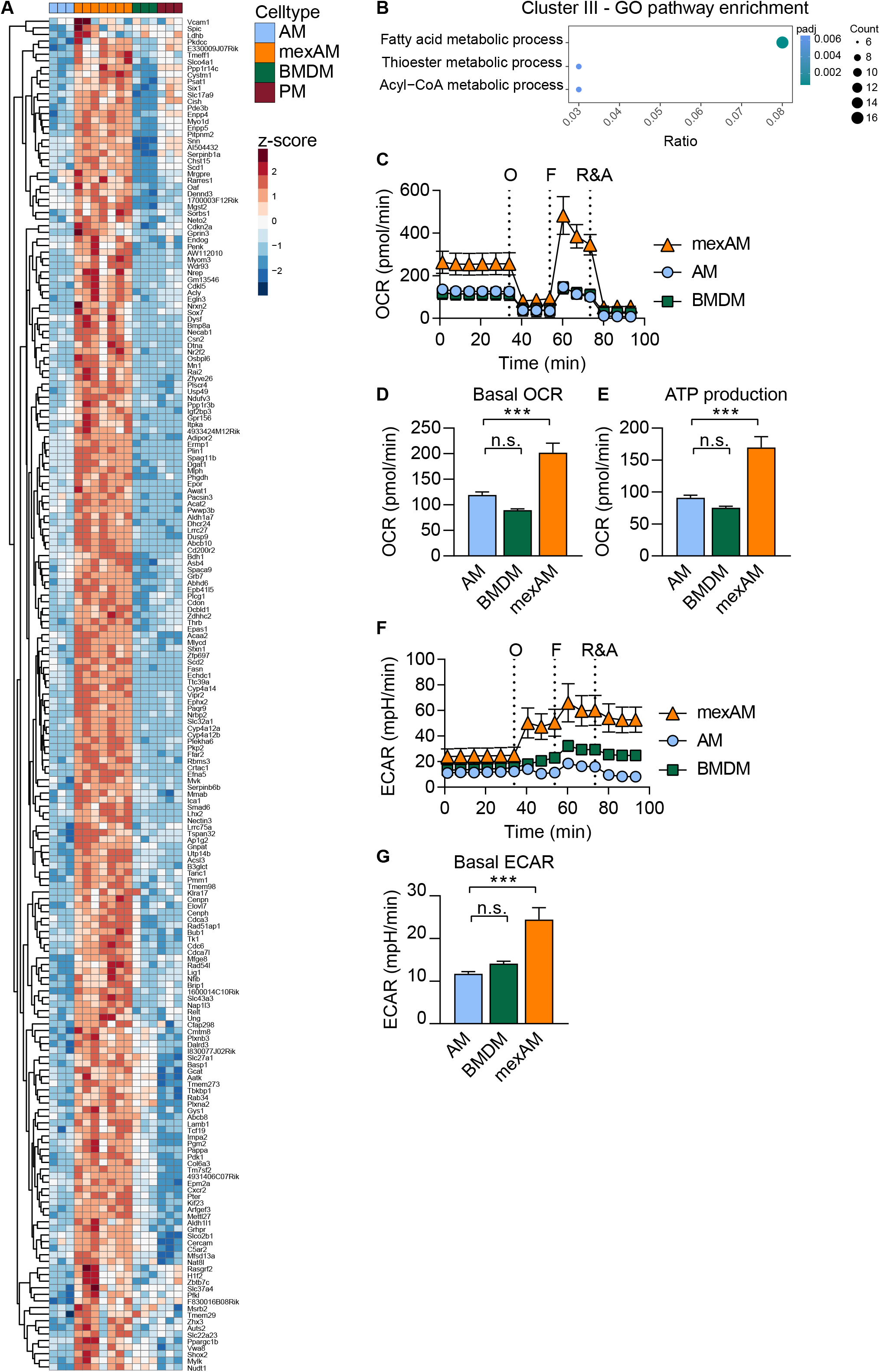
(**A**) Heatmap of genes found in cluster III (Fig. 3E). Raw counts were rlog transformed, followed by z-score scaling. (**B**) GO pathway enrichment of cluster III. (**C**-**G**) Oxygen consumption rate (OCR, **C**) and extracellular acidification rate (ECAR, **F**) of mexAMs, primary AMs or BMDMs (d5) were measured by Seahorse extracellular flux analysis at baseline and after injection of oligomycin (O), FCCP (F) and rotenone/antimycin A (R&A). Basal OCR (**D**), ATP production (**E**) and basal ECAR (**G**) of indicated cell types. (**C**-**G**) data are representative of two independent experiments. *p < 0.05, **p<0.01, ***p<0.001 (one-way ANOVA followed by Dunnett’s multiple comparison test). AM= alveolar macrophages, BMDM= bone marrow-derived macrophages, ECAR= extracellular acidification rate, GO= gene ontology, mexAM= mouse ex vivo cultured alveolar macrophages, ns= non significant, OCR= oxygen consumption rate, PM= peritoneal macrophages.

**Fig. S4.**
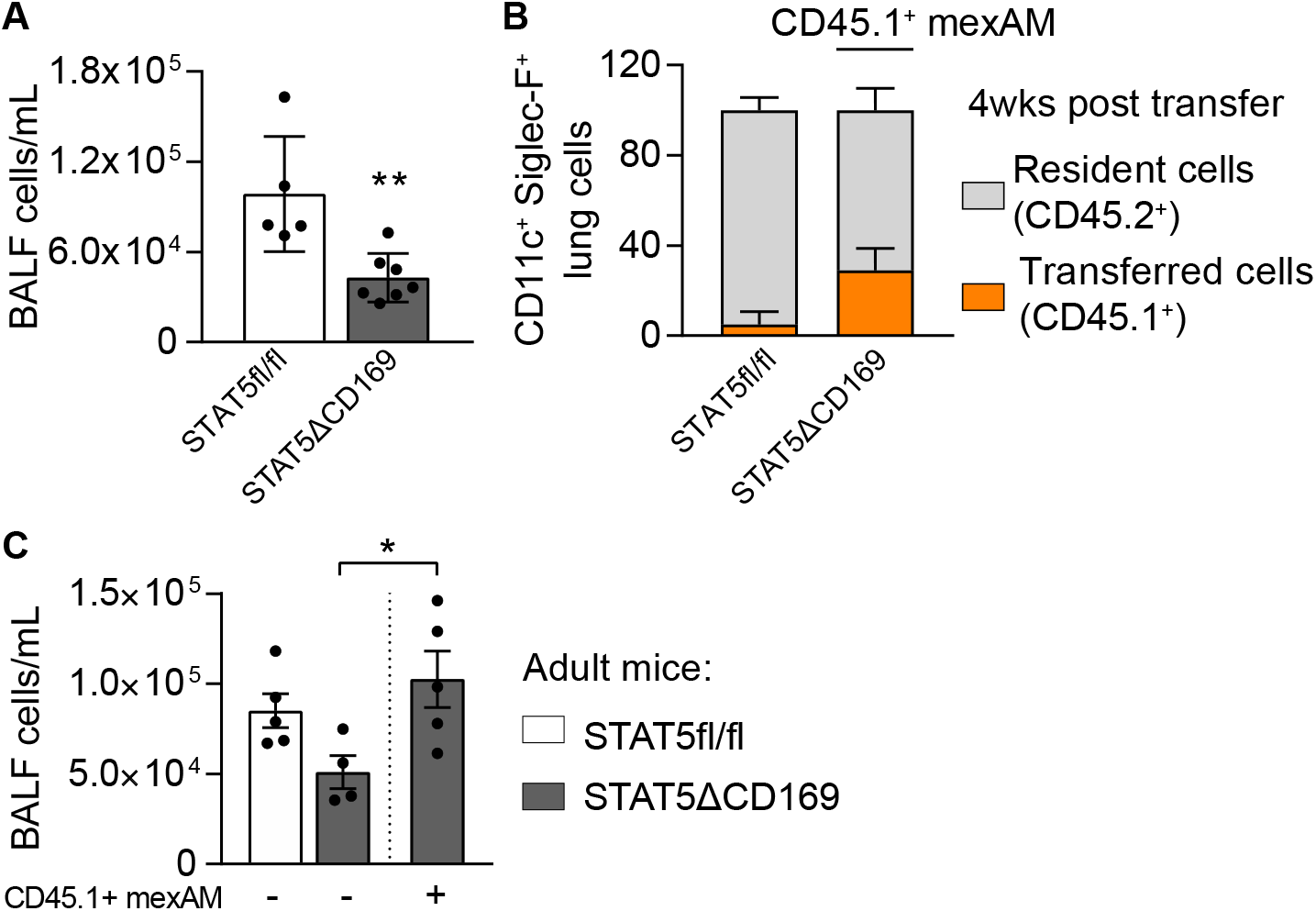
(**A**) BALF cell count per ml in STAT5fl/fl (control) and STAT5ΔCD169 mice. (**B**) Percentage of resident (CD45.2^+^, grey) and transferred (CD45.1^+^, orange) cells in lungs of control (STAT5fl/fl, n=4) and STAT5ΔCD169 mice (n=4) 4 weeks post CD45.1^+^ mexAM transfer. (**C**) BALF cell count per ml in STAT5fl/fl and STAT5ΔCD169 mice 8 weeks post transfer of CD45.1^+^ mexAMs. Graphs show means ± SEM of 4-5 biological replicates. *p < 0.05, **p<0.01 (Student’s t test (A) or one-way ANOVA followed by Sidak’s multiple comparison test (C)). BALF= bronchoalveolar lavage, mexAM= mouse ex vivo cultured alveolar macrophages.

**Fig. S5.**
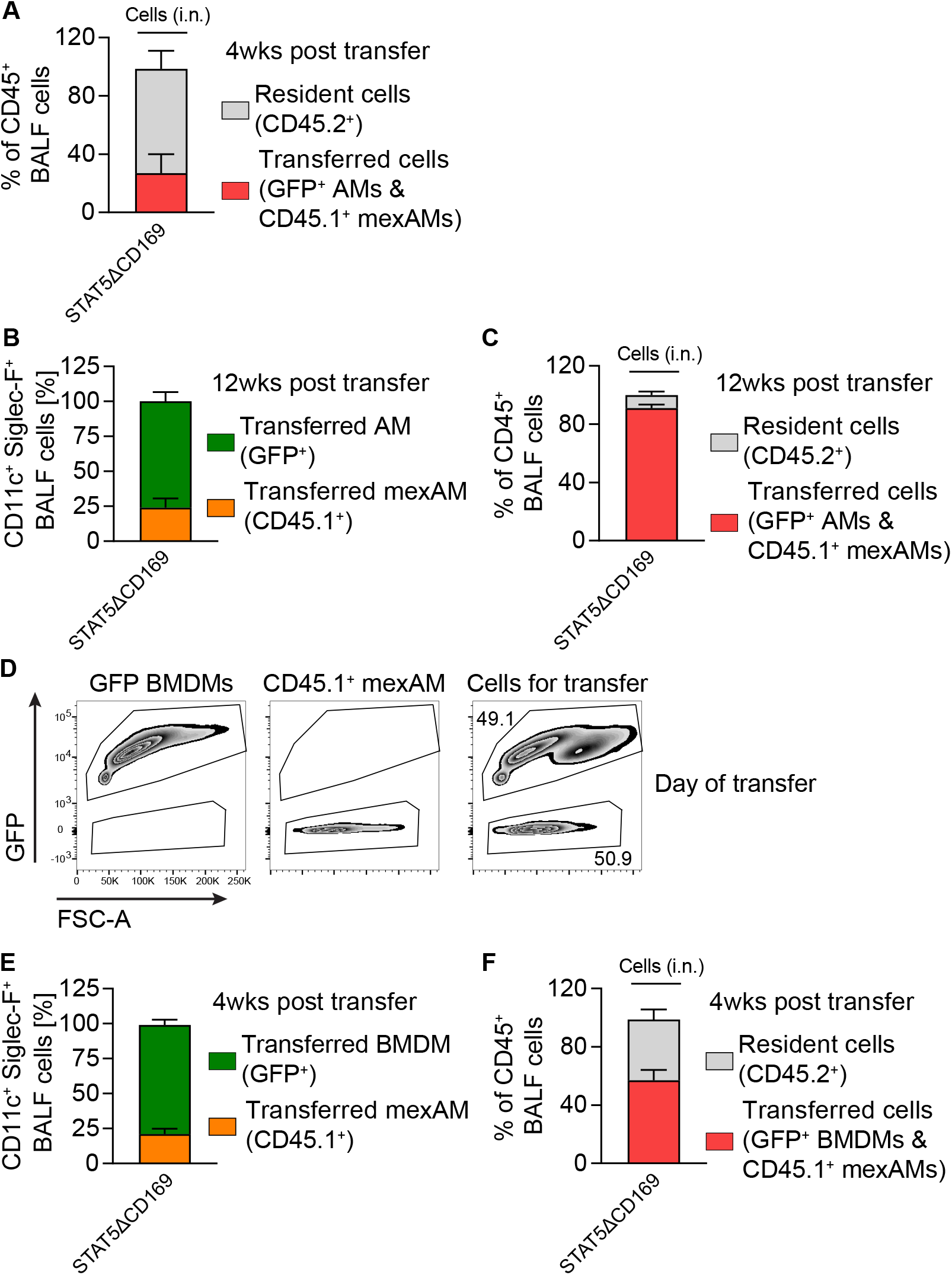
(**A**) Percentage of resident (CD45.2^+^, grey) and transferred (CD45.1^+^ mexAMs and GFP^+^ AMs, red) cells in BALF of STAT5ΔCD169 mice (n=4) 4 weeks post transfer. (**B**) Percentage of AMs (GFP^+^, green) and mexAMs (CD45.1^+^, orange) Siglec-F^+^CD11c^+^ cells in BALF of STAT5ΔCD169 (n=3) mice 12 weeks post transfer. (**C**) Percentage of resident (CD45.2^+^, grey) and transferred (CD45.1^+^ mexAMs and GFP^+^ AMs, red) cells in BALF of STAT5ΔCD169 mice (n=3) 12 weeks post transfer. (**D**) FACS analysis of GFP^+^ BMDM and CD45.1^+^ mexAM ratio on day of transfer. (**E**) BMDM (GFP^+^, green) and mexAM (CD45.1^+^, orange) percentage of Siglec-F^+^/CD11c^+^ cells in BALF of STAT5ΔCD169 (n=4) mice 4 weeks post transfer. (**F**) Percentage of resident (CD45.2^+^, grey) and transferred (CD45.1^+^ mexAMs and GFP^+^ BMDMs, red) cells in BALF of STAT5ΔCD169 mice (n=4) 4 weeks post transfer. Graphs show means ± SEM of 3-4 biological replicates. AM= alveolar macrophages, BALF= bronchoalveolar lavage, BMDM= bone marrow-derived macrophages, i.n.= intranasal, mexAM= mouse ex vivo cultured alveolar macrophages.

## Supplemental Tables

Resource table

Table S1 (Legendplex data for AM, mexAM and BMDM samples, related to Figure 2)

Table S2 (DEGs rlog transformed including cluster annotation and gene symbol, related to Figure 3)

Table S3 (List of AM specific genes, related to Figure 3)

